# The Prevalence and Impact of Model Violations in Phylogenetics Analysis

**DOI:** 10.1101/460121

**Authors:** Suha Naser-Khdour, Bui Quang Minh, Wenqi Zhang, Eric Stone, Robert Lanfear

## Abstract

In phylogenetic inference we commonly use models of substitution which assume that sequence evolution is stationary, reversible and homogeneous (SRH). Although the use of such models is often criticized, the extent of SRH violations and their effects on phylogenetic inference of tree topologies and edge lengths are not well understood. Here, we introduce and apply the maximal matched-pairs tests of homogeneity to assess the scale and impact of SRH model violations on 3,572 partitions from 35 published phylogenetic datasets. We show that many partitions (39.5%) reject the SRH assumptions, and that for most datasets, the topologies of trees inferred from all partitions differ significantly from those inferred using the subset of partitions that do not reject the SRH assumptions. These results suggest that the extent and effects of model violation in phylogenetics may be substantial. They highlight the importance of testing for model violations and possibly excluding partitions that violate models prior to tree reconstruction. They also suggest that further effort in developing models that do not require SRH assumptions could lead to large improvements in the accuracy of phylogenomic inference. The scripts necessary to perform the analysis are available in https://github.com/roblanf/SRHtests, and the new tests we describe are available as a new option in IQ-TREE (http://www.iqtree.org).

## Introduction

Phylogenetics is an essential tool for inferring evolutionary relationships between individuals, species, genes, and genomes. Moreover, phylogenetic trees form the basis of a huge range of other inferences in evolutionary biology, from gene function prediction to drug development and forensics (Eisen 1998; Farrell, et al. 2000; Mäser, et al. 2001; Gardner, et al. 2002; Yao, et al. 2003; Grenfell, et al. 2004; Yao, et al. 2004; Salipante and Horwitz 2006; Gray, et al. 2009; Brady and Salzberg 2011; Dunn, et al. 2011).

Most phylogenetic studies use models of sequence evolution which assume that the evolutionary process follows stationary, reversible and homogeneous (SRH) conditions. Stationarity implies that the marginal frequencies of the nucleotides or amino acids are constant over time, reversibility implies that the evolutionary process is stationary and undirected, and homogeneity implies that the instantaneous substitution rates are constant along the tree or over an edge (Felsenstein 2004; Yang and Rannala 2012; Jermiin, et al. 2017). However, these simplifying assumptions are often violated by real data (Foster and Hickey 1999; Tarrío, et al. 2001; Paton, et al. 2002; Goremykin and Hellwig 2005; Murray, et al. 2005; Bourlat, et al. 2006; Hyman, et al. 2007; Sheffield, et al. 2009; Nesnidal, et al. 2010; Nabholz, et al. 2011; Martijn, et al. 2018). Such model violation may lead to systematic error that, unlike stochastic error, cannot be solved simply by increasing the size of a dataset (Felsenstein 2004; Ho and Jermiin 2004; Jermiin, et al. 2004; Philippe, et al. 2005; Sullivan and Joyce 2005; Kumar, et al. 2012; Brown and Thomson 2017; Duchene, et al. 2017). As phylogenetic datasets are steadily growing in terms of taxonomic and site sampling, it is vital that we develop and employ methods to measure and understand the extent to which systematic error affects phylogenetic inference (systematic bias), and explore ways of mitigating systematic bias in empirical studies.

One approach to accommodate data that have evolved under non-SRH conditions is to employ models that relax the SRH assumptions. A number of non-SRH models have been implemented in a variety of software packages (Foster 2004; Lartillot and Philippe 2004; Blanquart and Lartillot 2006; Boussau and Gouy 2006; Jayaswal, et al. 2007; Knight, et al. 2007; Dutheil and Boussau 2008; Jayaswal, et al. 2011; Sumner, et al. 2012; Zou, et al. 2012; Groussin, et al. 2013; Jayaswal, et al. 2014; Nguyen, et al. 2015; Woodhams, et al. 2015). However, such models remain infrequently used, as searching for optimal phylogenetic trees under these models is computationally demanding (Betancur-r, et al. 2013) and the implementations are often not easy to use. As a result, the vast majority of empirical phylogenetic inferences rely on models that assume sequences have evolved under SRH conditions, such as the general time reversible (GTR) family of models implemented in the most widely-used phylogenetics software packages (Swofford 2001; Drummond and Rambaut 2007; Guindon, et al. 2010; Ronquist, et al. 2012; Bazinet, et al. 2014; Bouckaert, et al. 2014; Stamatakis 2014; Nguyen, et al. 2015; Höhna, et al. 2016).

Another approach to accounting for data that may have evolved under non-SRH conditions is to test for model violations prior to tree reconstruction. Here, one first screens datasets or parts of datasets, and reconstruct trees exclusively from data that do not reject SRH conditions. A number of methods have been proposed to test for violation of SRH conditions in aligned sequences prior to estimating trees (Bowker 1948; Stuart 1955; Rzhetsky and Nei 1995; Kumar and Gadagkar 2001; Weiss and von Haeseler 2003; Ababneh, et al. 2006; Ho, et al. 2006), and there are also *a posteriori* tests for absolute model adequacy which are employed after trees have been estimated (Goldman 1993; Brown and ElDabaje 2009; Brown 2014; Duchene, et al. 2017; Brown and Thomson 2018).

Allowing the data to reject the model when the assumptions of the model are violated is an important approach to reducing systematic bias in phylogenetic inference (Philippe, et al. 2005; Brown 2014). Knowing in advance which sequences and loci are inconsistent with the SRH assumptions will allow us to choose more complex models for phylogeny reconstruction of this data or to omit some of these sequences and loci from downstream analyses (Kumar and Gadagkar 2001). The need for methods that assess the evolutionary process prior to phylogenetic inference becomes more important as the number of sequences and sites per dataset increases, because systematic bias has an increasing effect on inferences from larger phylogenetic datasets (Ho and Jermiin 2004; Jermiin, et al. 2004; Phillips, et al. 2004; Delsuc, et al. 2005).

In this paper we evaluate the extent and effect of model violation due to non-SRH evolution using 35 empirical datasets with a total of 3,572 partitions. We determine if the SRH assumptions are violated by extending and applying the matched-pairs tests of homogeneity (Jermiin, et al. 2017) to each partition. We then compare the phylogenetic trees for each dataset estimated from all of the partitions, the partitions that reject the SRH assumptions, and the partitions that do not reject the SRH assumptions, in order to evaluate the effect violating SRH conditions on phylogenetic inference.

## Materials and Methods

### Empirical datasets

In order to assess the impact of model violation in phylogenetics, we first gathered a representative sample of 35 partitioned empirical datasets that had been used for phylogenetic analysis in recent studies (Table 1). Within the constraints of selecting data that were publicly available and suitably annotated, i.e. such that all loci and all codon positions within protein-coding loci could be identified, we selected the datasets to provide as representative a sample as possible of the data types, taxa, and genomic regions most commonly used to infer bifurcating phylogenetic trees from concatenated alignments. These datasets include nucleotide sequences from nuclear, mitochondrial, plastid and virus genomes, and include protein-coding DNA, introns, intergenic spacers, tRNA, rRNA and ultra-conserved elements. The number of taxa and sites in these datasets range from 27 to 355 and from 699 to 1,079,052 respectively. The clades represented in these datasets include animals, plants and viruses. We partitioned all datasets to the maximum possible extent based on the biological properties of the data, i.e. we divided every locus and every codon position within each protein-coding locus into a separate partition. All partitioning information is available at the github repository https://github.com/roblanf/SRHtests/tree/master/datasets, and the full details of every dataset are provided in Table 1 and in extended Table 11.

**Table 1.**
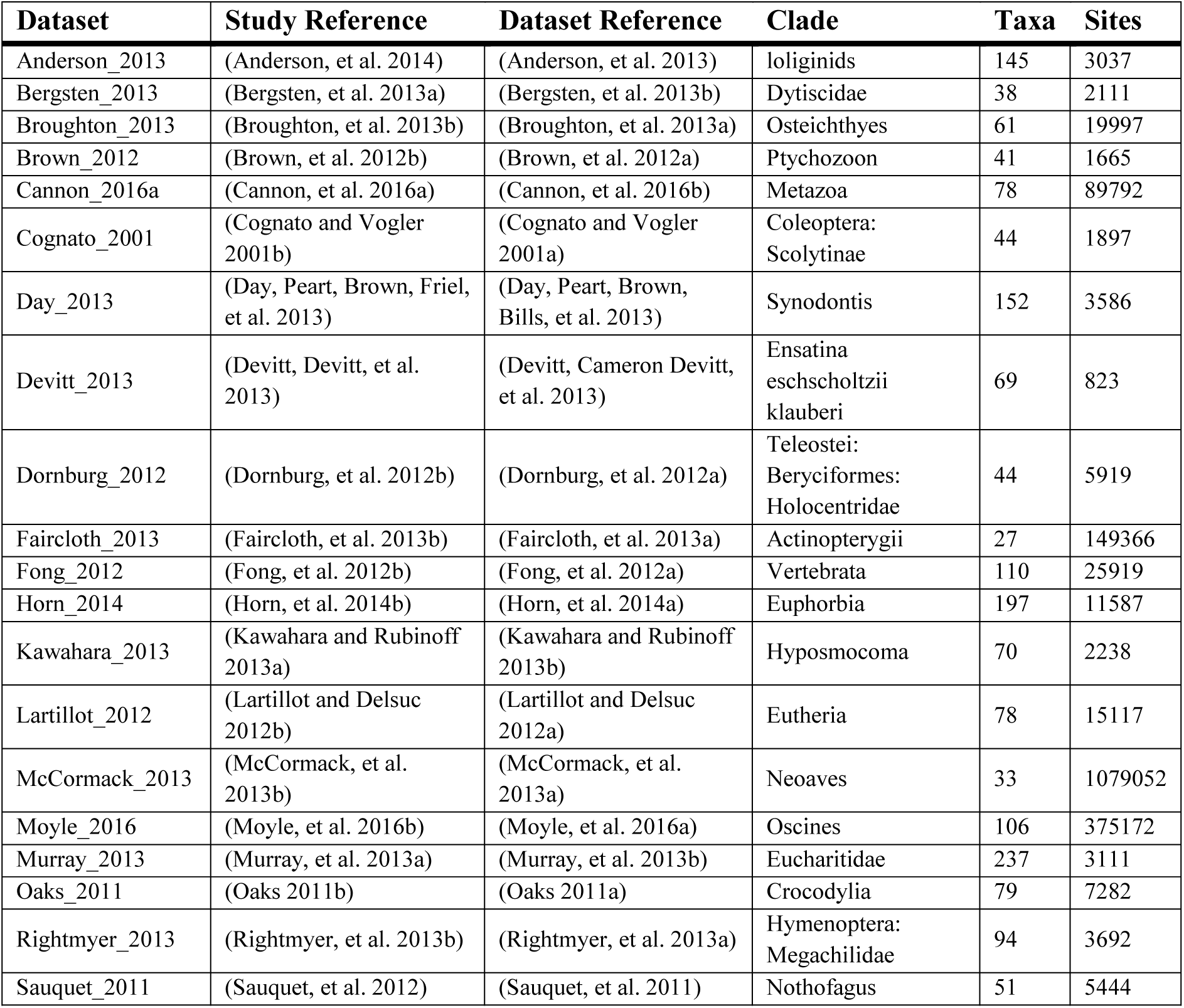
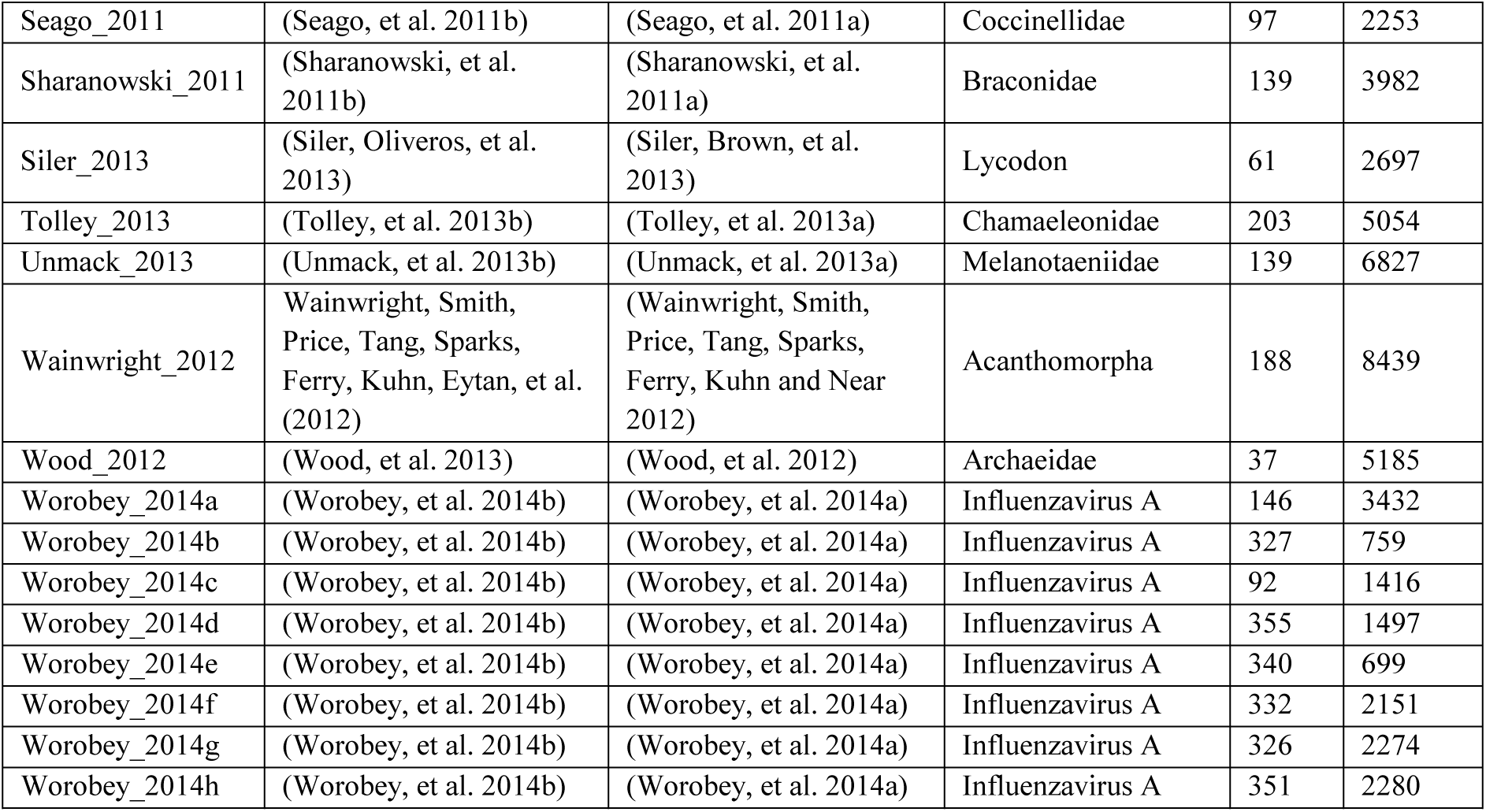
Number of taxa, number of sites, clade and study reference for each dataset that has been used in this study

### Workflow summary

Figure 1 outlines the workflow. For each partition in each dataset, we used two approaches based on the three matched-pairs tests of homogeneity to ask whether the evolution of the aligned sequences in the partition rejects the SRH assumptions. The three matched-pairs tests of homogeneity, described in more detail below, test three slightly different assumptions about the historical process that generated each aligned pair of sequences in a given partition. A significant result from any test suggests that the nature of the evolutionary process required to explain the aligned sequences violates at least one of the three SRH conditions (Jermiin, et al. 2017). For each test, we classify each partition as *pass* if the result of the test is non-significant or *fail* if the result of the test is significant. We then denote the original dataset as D_all_, while the concatenation of *pass* partitions is denoted D_pass_ and the concatenation of *fail* partitions as D_fail_ (Fig. 1).

**Fig. 1.**
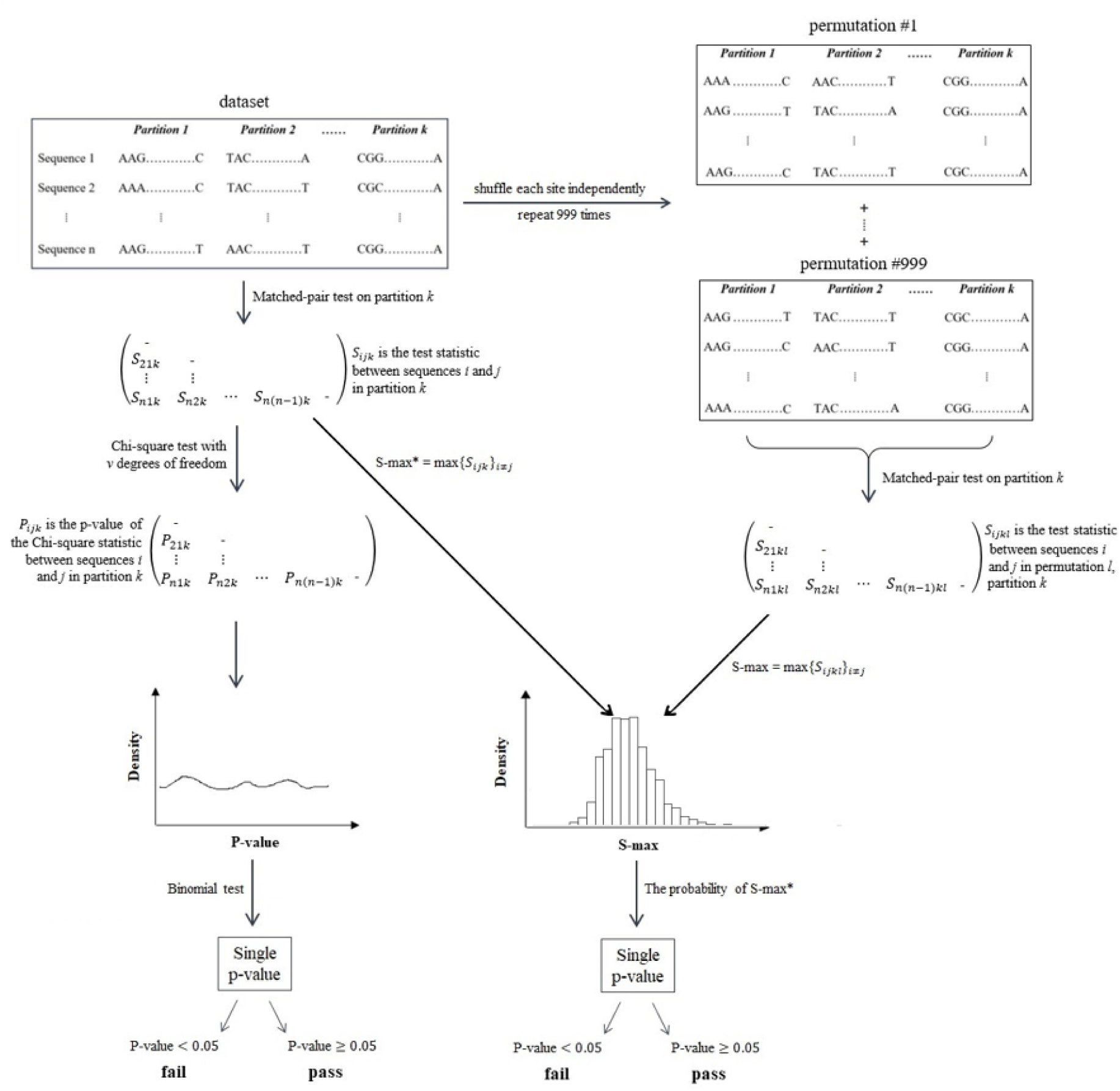
Flow chart of methodology. After application of the matched-pairs tests of homogeneity on each possible pair of sequences for each partition, we have two options: 1) apply a binomial test on the p-values of all possible pairs of sequences of each partition to derive a P-value for that partition. 2) Take the maximum test statistic value and compare it to a null distribution of the maximum test statistics derived from permutation of the sites of the alignments.

To investigate the impact of model violation on phylogenetic inference, we infer and compare three phylogenetic trees, T_all_, T_pass_ and T_fail_, estimated from D_all_, D_pass_ and D_fail_, respectively.

### Matched-pairs tests of homogeneity

The three matched-pairs of homogeneity that are applied to pairs of sequences are: the MPTS (matched-pairs test of symmetry), MPTMS (matched-pairs test of marginal symmetry), and MPTIS (matched-pairs test of internal symmetry). The statistics are computed on a *m*-by-*m* (*m* is 4 for nucleotides and 20 for amino acids) divergence matrix *D* with elements *d*_*ij*_, where *d*_*ij*_ is the number of alignment sites having nucleotide (or amino acid) *i* in the first sequence and nucleotide (or amino acid) *j* in the second sequence.

The MPTS tests the symmetry of D by computing the Bowker’s test statistic as the chi-square distance between D and its transpose:

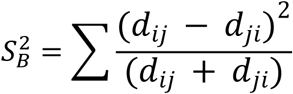

A p-value is then obtained by a chi-square test with *m*(*m* − 1)/2 degrees of freedom. A low p-value (<0.05) indicates that the assumption of symmetry is rejected and evolution is non-stationary or non-homogeneous (Jermiin, et al. 2017).

The MPTMS tests the equality of nucleotide or amino acid composition between two sequences. To do so, MPTMS computes the Stuart’s test statistic based on the difference between nucleotide or amino acid frequencies of two sequences, *u*, and its variance-covariance matrix, 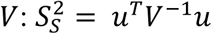.

Where *u* is the vector of marginal differences and *u*^^*T*^^ = (*d*_1•_ − *d*_•1_, *d*_2•_ − *d*_•2_, …, *d*_*k*•_ − *d*_•*k*_). *d*_*i*•_ is the sum of *d*_*ij*_ over *j*, *d*_•*j*_ is the sum of *d*_*ij*_ over *i*, and *k* = *m* − 1.

*V* is the estimated variance-covariance matrix of *u* under the assumption of marginal symmetry with the elements

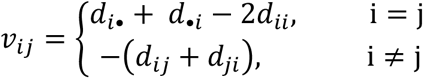

A p-value is obtained by a chi-square test with *m* − 1 degrees of freedom. A low p-value (<0.05) indicates that the stationarity assumption is rejected.

The MPTIS uses the test statistic as the difference between Bowker’s and Stuart’s statistic: 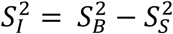. Hence, it is also chi-square distributed and one obtains a p-value with (*m* − 1)(*m* − 2)/2 degrees of freedom. A low p-value (<0.05) indicates that the homogeneity assumption is rejected.

The MPTS, MPTMS and MPTIS test different aspects of the symmetry with which substitutions accumulate between pairs of sequences: The MPTS is a comprehensive and sufficient test to determine whether the data complies with the SRH assumptions (Jermiin, et al. 2017), but it cannot provide any information about the source of this violation. Some information on the underlying source of model violation may be obtained by performing the other two tests of symmetry, the MPTMS and the MPTIS. If the violation of the SRH assumptions stems from differences in base composition between the sequences, this should affect the marginal symmetry of the sequence pair, which can in principle be detected by the MPTMS. While if the violation of the SRH assumptions stems from differences in substitution rates over time, this should affect the internal symmetry of the sequence pair, which can in principle be detected by the MPTIS. However, even after performing all three tests, it is difficult to ascertain which of the three SRH assumptions is violated during the evolutionary process because the relationships between the SRH conditions and the three matched-pair tests is neither bijective nor injective, i.e. there is no one-to-one correspondence between the three tests and violation of the three SRH conditions (Jermiin, et al. 2017). However, the three matched-pairs tests of homogeneity were designed to ask whether any single pair of sequences rejects the SRH conditions (Jermiin, et al. 2017). To ask whether a given partition rejects SRH conditions, we developed two approaches to extend the matched-pairs tests of homogeneity to accommodate datasets with more than two sequences.

### Extending the matched-pairs tests of homogeneity to multiple sequence alignments

There are many potential ways to extend the three matched pairs tests of homogeneity for multiple sequence alignments. One approach is to compute the p-value for all pairs of sequences in an alignment, and then ask whether the distribution of the resulting p-values follows the distribution expected under the null hypothesis. In this approach, we apply the matched-pair tests of homogeneity to every pair of sequences in an alignment, resulting in 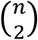 chi-square p-values for each partition, where *n* is the number of sequences in the partition (fig. 1). Under the null hypothesis of SRH evolution, the marginal distribution of each p-value should be uniform on the interval [0,1], suggesting a 5% chance that any one of them falls below 0.05. In principle, we could use this logic to assess whether we observe smaller p-values than we would expect by chance, in which case a partition would be deemed to fail the SRH conditions. Specifically, we could count how many chi-square p-values are less than 0.05 and compare the result to a Binomial distribution with 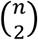 trials and success probability 0.05. If the corresponding Binomial p-value were smaller than 0.05, we would classify the partition as *fail*; otherwise, the partition is labelled as *pass*.

Despite its appealing simplicity, this approach suffers from the serious drawback that it ignores the dependencies among p-values. P-values are dependent because many pairs of sequences will cross shared branches in the tree. Thus, a full accounting of the dependencies among p-values would require knowledge of the underlying phylogeny. Given that these tests aim to determine *a priori* whether it is feasible to build a reliable phylogeny, it would be inappropriate to use an estimated phylogeny as part of the test. Because of this limitation, we only present the results of this test in the supplementary material. To avoid confusion between the binomial and pairwise tests, we denote the binomial extensions of the MPTS, MPTMS, and the MTPIS as BiSymTest, BiSymTest_mar_, and BiSymTest_int_ respectively.

### Maximum statistic approach

The second approach, which we call *MaxSymTest*, to determine whether a given alignment rejects SRH conditions, is to consider only the pair of taxa with the maximum test statistic value (which we denote as *S*_*max*_). MaxSymTest assumes that model violations, if present, would occur along the path connecting these two taxa. MaxSymTest overcomes the non-independence issue because it uses data from just a single pair of sequences from an alignment. Furthermore, it does not require the knowledge of the underlying tree topology. Because we do not assume any distribution of *S*_*max*_, we assess its statistical significance as follows. We compute the null distribution for *S*_*max*_ by permutating the sequences for each alignment site independently. Specifically, for a single test of an *S*_*max*_ value, we generate 999 permutated alignments (Fig. 1) and use these to calculate the corresponding null distribution comprised of 999 *S*_*max*_ values. MaxSymTest then assigns a p-value as the fraction of permutated *S*_*max*_ larger than or equal to the *S*_*max*_ from the original alignment. For convenience, we denote the maximum value approach of the MPTS, MPTMS, and MPTIS as MaxSymTest, MaxSymTest_mar_, and MaxSymTest_int_ respectively. We present the results of the MaxSymTest analyses in the main text. The supplementary information contains a side-by-side comparison of the MaxSymTest and the BiSymTest results.

### Phylogenetic inference

We used IQ-TREE (Nguyen, et al. 2015) to infer up to seven phylogenetic trees for every dataset: T_all_ (all partitions from the original dataset; D_all_); and T_pass_ and T_fail_ based on the D_pass_ and D_fail_ datasets from each of the three tests (MaxSymTest, MaxSymTest_mar_, MaxSymTest_int_), provided that there was at least one partition in each category. We ran IQ-TREE using the default settings with the best-fit fully-partitioned model (Chernomor, et al. 2016), which allows each partition to have its own evolutionary model and edge-linked rate determined by ModelFinder (Kalyaanamoorthy, et al. 2017) followed 1000 ultrafast bootstrap replicates (Hoang, et al. 2018).

### Distance between trees

For each of the three tests (MPTS, MPTMS, MPTIS) we calculated the Normalised Path-Difference (NPD) and quartet distance (QD) (Steel and Penny 1993; Sand, et al. 2014) between all three possible pairs of trees (T_all_ vs. T_pass_; T_all_ vs. T_fail_; and T_pass_ vs. T_fail_), as long as D_pass_ and D_fail_ were non-empty and so T_pass_ and T_fail_ had been estimated. The path-difference metric (PD) is defined as the Euclidean distance between pairs of taxa (Steel and Penny 1993; Mir and Russello 2010). In this study, because we are interested only in differences between topologies, we use the variant of the PD metric that ignores branch lengths. In order to compare path distances between trees with different number of taxa, we normalised PD (to obtain NPD) by the mean of a null distribution of PDs generated from 10K random pairs of trees with the same number of taxa (Bogdanowicz, et al. 2012). Thus, an NPD of zero indicates an identical pair of trees, an NPD of 1 indicates that a pair of trees is as similar as a pair of randomly-selected trees with the same number of taxa; and an NPD greater than 1 indicates a pair of trees that are less similar than a randomly-selected pair of trees with the same number of taxa. Since path differences are always non-negative, the NPD is also guaranteed to be non-negative.

The QD metric is defined as the fraction of quartets (subsets of four taxa) that induce different subtrees between two comparing trees. QD ranges between 0 and 1, where 0 means that two trees are identical and 1 means that they do not share any quartet subtrees. Compared with PD, QD has a main advantage that its distribution is less sensitive to the underlying distribution on tree topologies (Steel and Penny 1993).

### Tree topology tests

The NPD and the QD give us measures of the differences between pairs of trees, but they do not tell us whether the differences are phylogenetically significant in the three datatsets (D_pass_, D_all_, and D_fail_) derived from a given test. For example, trees that differ due to stochastic error _associated with small datasets may be very different, but such differences may not be_ statistically significant. To assess the significance of the differences between T_pass_, T_all_ and T_fail_, we used the weighted Shimodaira-Hasegawa (wSH) test (Shimodaira and Hasegawa 1999; Shimodaira 2002) implemented in IQ-TREE with 1000 RELL replicates (Kishino, et al. 1990). Given the alignment (D_pass_), the wSH test computes a p-value for each tree, where a low p-value (<0.05) implies that the corresponding tree has a significantly worse likelihood than the best tree in the set of T_pass_, T_all_ and T_fail_. We use D_pass_ for these tests because it is, by definition, the only dataset that does not reject the underlying assumptions of the SH test. As such, we can only compute sWH p-values when D_pass_ is non-empty. Thus, we performed two sWH tests for each of the three MaxSymTest variant: one that asks whether T_all_ can be rejected in favour of T_pass_, and another that asks whether T_fail_ can be rejected in favour of T_pass_. In cases where there were no partitions in D_pass_ or D_fail_, we were unable to perform the wSH test.

### Correlation between number of substitutions and model violation

We hypothesised that partitions with more substitutions may be more likely to violate the SRH assumptions, since substitutions form the raw data for the matched-pairs tests of symmetry. To assess this, we fitted a linear mixed-effects model for each of the three tests using the glmer function from the lme4 package in R (Bates, et al. 2015). In this model, we treat each partition as a datapoint, the number of substitutions measured for that partition as a fixed effect, and the dataset from which that partition was taken as a random effect. This allows us to estimate the extent to which the number of substitutions in a partition correlates with whether a partition fails a given test of symmetry, after accounting for differences between the datasets. To calculate the R squared value we use the r.squaredGLMM function from the MuMIn package in R (Barton 2009; Nakagawa and Schielzeth 2013).

### Software implementation

We implemented a new option --maxsymtest NUM (where NUM specifies the number of permutations) in IQ-TREE to perform the three MaxSymTest matched pairs tests of symmetry. In addition, the option --symtest-remove-bad allows users to remove from the final analysis partitions that fail the MaxSymTest. One can change the removal criterion to MaxSymTest_mar_ or MaxSymTest_int_ via the --symtest-type MAR|INT option. In addition, the cut-off p-value can be changed using the --symtest-pval NUM option, where the default value is 0.05.

### Reproducibility

The GitHub repository https://github.com/roblanf/SRHtests contains the raw data and Python and R scripts necessary to perform all analyses reported in this study.

## Results

### Violation of SRH conditions is common across 35 empirical datasets

Across all 3,572 partitions analysed, 1,475 (40.8%) failed the MaxSymTest, 1,483 (41.5%) failed the MaxSymTest_mar_, and 312 (8.7%) failed the MaxSymTest_int_. 1,804 (50.5%) of partitions failed at least one test.

The proportion of partitions failing each test varied substantially among datasets (Fig. 2), but on average 44.9% of the partitions in each dataset failed the MaxSymTest, 41.8% failed the MaxSymTest_mar_, and 8.2% failed the MaxSymTest_int_.

**Figure 2.**
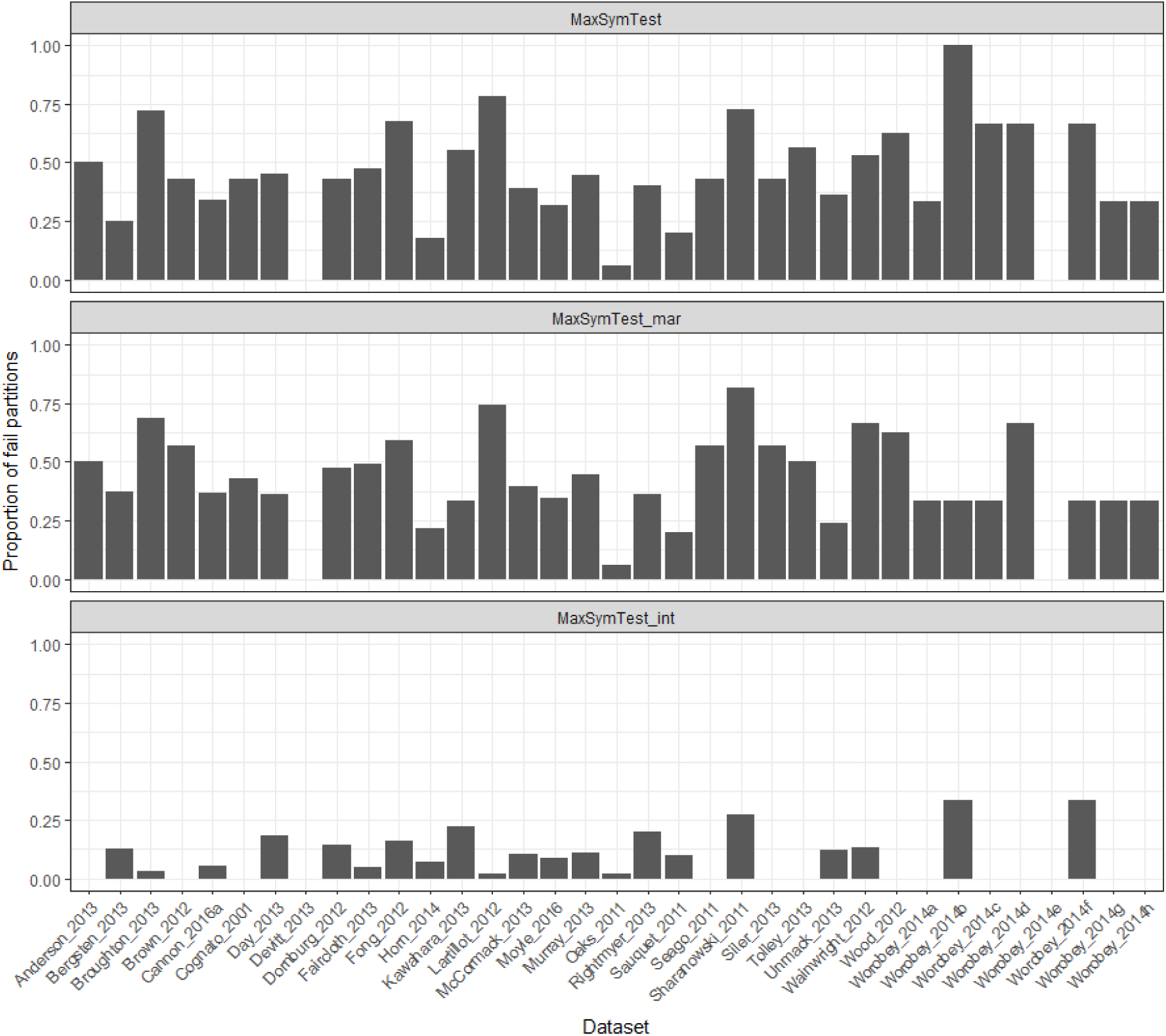
The proportion of partitions that reject the null hypothesis of the MaxSymTest, MaxSymTest_mar_ and MaxSymTest_int_ (p-value < 0.05) in each dataset. The fraction of failing partitions also varied with the type of genome (e.g. mitochondrial, chloroplast, or nuclear) and the type of locus (e.g. protein-coding, UCE, tRNA) from which the partition was sequenced (Table 2) although we note that a substantial proportion of the partitions from almost every category failed at least one of the tests (Table 2). Similar results are detected with the BiSymTest (Extended Table 1)

**Table 2.** The proportion of partitions that failed at least one of the three tests - MaxSymTest, MaxSymTest_mar_, MaxSymTest_int_.

| Type / genome | nuclear | mitochondrial | plastid | virus |
| --- | --- | --- | --- | --- |
| 1<sup>st</sup> codon positions | 51.1% | 53.3% | 16.7% | 12.5% |
| 2<sup>nd</sup> codon positions | 48.0% | 36.7% | 50% | 37.5% |
| 3<sup>rd</sup> codon positions | 83.6% | 70.0% | 16.7% | 87.5% |
| Other (e.g. intron) | 27.8% | 66.7% | 16.7% |  |
| rRNA | 70.0% | 25.0% |  |  |
| UCE | 49.3% |  |  |  |
| tRNA |  | 12.5% |  |  |

There were no clear differences in the substitution models that were selected for the partitions that pass or fail the tests (see Extended Tables 2-6). However, we note that the two most-frequently selected substitution models (for >35% of the partitions) were relatively simple: K80 (Kimura 1980) and HKY (Hasegawa, et al. 1985). K80 has one single parameter, the transition to transversion rate ratio. Whereas HKY additionally has four base frequency parameters.

### Model violation has a large influence on tree topologies

For MaxSymTest and MaxSymTest_mar_ and according to two different tree distance metrics (NPD and QD), we find that the tree inferred from the original dataset (T_all_) was more similar to the tree estimated from the failed partitions T_fail_ (Table 3, Extended tables 7, 9-10) compared with T_pass_. Furthermore, the mean NPD distance between T_pass_ and T_fail_ across all 35 datasets for the MaxSymTest was 0.71, i.e., they are 71% as dissimilar as random pairs of trees. This suggests that violations of SRH assumptions drive changes in tree topologies. The results of the wSH tests (Table 4, Extended Table 8) confirm that the differences between trees that we observe tend to be statistically significant. For example, when using the MaxSymTest, T_pass_ rejects T_all_ in ~48% of the datasets, and T_fail_ in ~84% of the datasets.

**Table 3.** The proportion of datasets that have the highest NPD metric and QD metric respectively between the three comparisons (all-fail, all-pass, pass-fail) for MaxSymTest, MaxSymTest_mar_, and MaxSymTest_int_.

| MaxSymTest |  | T<sub>fail</sub> | T<sub>pass</sub> |
| --- | --- | --- | --- |
|  | T<sub>all</sub> | 0.0% (0.0%) | 35.0% (4.8%) |
|  | T<sub>pass</sub> | 65.0% (95.2%) |  |
| MaxSymTest<sub>mar</sub> | T<sub>all</sub> | 0.0% (0.0%) | 24.0% (4.0%) |
|  | T<sub>pass</sub> | 76.0% (96.0%) |  |
| MaxSymTest<sub>int</sub> | T<sub>all</sub> | 12.5% (12.5%) | 0.0% (0.0%) |
|  | T<sub>pass</sub> | 87.5% (87.5%) |  |

**Table 4.** The proportion of datasets that have a significant p-value in the weighted SH test when using D_pass_ as the input alignment for the test.

|  | T<sub>all</sub> | T<sub>fail</sub> |
| --- | --- | --- |
| MaxSymTest | 48% | 84% |
| MaxSymTest<sub>mar</sub> | 50% | 82% |
| MaxSymTest<sub>int</sub> | 8% | 44% |

### The number of substitutions explains less than fifth of the variance in passing or failing the tests of symmetry

The number of substitutions in a partition explained 17% of the variation in whether or not a partition passed or failed the MaxSymTest (Extended figs. 6-7). This proportion is very similar for MaxSymTest_mar_ (18%) (Extended figs. 8-9), but is dramatically lower for the MaxSymTest_int_ (2%) (Extended figs. 10-11). Accordingly, the number of substitutions in a partition is a highlight significant (p<2e-16) predictor of passing or failing any of the tests. However, that the number of substitutions explains less than a fifth of the variation suggests that other factors, for example underlying differences in the extent to which partitions violate the SRH assumptions, are driving the remaining ~80% of the variation.

### Model violation affects the internal relationships of Spiralia and the position of Xenacoelomorpha

To examine the effects of model violation in more detail, we selected a single dataset for more detailed consideration. Conflicting support for the placement of Xenacoelomorpha, the clade that contains Xenoturbella and Acoelomorpha, in the tree of life across different analyses has led to various hypotheses regarding to the evolution of Bilateria (Cannon, et al. 2016a). It has been suggested that such inferences might be strongly affected by model violation and systematic error (Philippe, et al. 2011). To assess whether data that pass or fail the MaxSymTest show different signals regarding the evolution of the Bilateria, we examined in more detail the T_all_, T_pass_, and T_fail_ trees from a recent study that addressed this question (2016a). This dataset comprises 76 metazoan taxa, 2 choanoflagellate outgroups, 212 genes and 424 partitions representing the first and second codon positions of the 212 genes (Cannon, et al. 2016b). The tree reconstructed from all of the partitions (T_all_) is identical to the tree reconstructed from the partitions that pass the MaxSymTest (T_pass_), and both are identical to the tree shown in the original paper from both DNA and amino acid data (Canon, et al. 2016a), which places Xenacoelomorpha as the sister group of Nephrozoa (Deuterostomia and Protostomia) with 100% bootstrap support (Fig. 4, Extended figs. 1-3).

The tree reconstructed from the data that fail the MaxSymTest (T_fail_) on the other hand (inferred from 143 partitions) shows Xenoturbella as the sister group to Nephrozoa with 93% bootstrap support (Fig. 4, Extended figs. 4-5). It also shows Platyzoa (Rotifera, Platyhelminthes and Gastrotricha) as a sister of Acoelomorpha with 95% bootstrap support. It is clear from the results that the partitions that comprise D_fail_ support a very different set of relationships to those that comprise D_pass_.

## Discussion

In this paper, we show that model violation is prevalent and has a strong impact on tree reconstruction in many phylogenetic datasets. This impact varies a lot between different datasets and different types of partitions. The trees inferred from different groups of partitions from the same dataset often have topologies that are biologically and statistically significantly different.

Our results show great variation in the extent of model violation among different datasets and partitions. This is demonstrated by the different proportion of partitions that failed the matched-pairs tests of symmetry in each dataset and in each genomic context (codon position, rRNA, tRNA, UCE or other) and type of genome (nuclear, mitochondrial, plastid and virus). Model violations are most frequently observed in the third codon positions for viral, mitochondrial and nuclear genomes and intergenic spacers in plastid sequences. Yet, our results affirm that non-SRH evolution is not constrained to these genomic regions. For example, in a dataset of first and second codon positions from 212 genes (Cannon, et al. 2016b), 34% of partitions showed significant evidence of violating the SRH assumptions according to the MaxSymTest. The tree inferred from the partitions that show significant model violation differs a great deal in its topology from the tree inferred from the partitions that do not show significant model violation, particularly with respect to the placement of the focal taxon Xenoturbella (fig. 3). From looking at the results of the two other tests – MaxSymTest_mar_ and MaxSymTest_int_, we noticed that all the partitions that failed the MaxSymTest also failed the MaxSymTest_mar_, suggesting that those partitions are violating the models mainly due to non-stationarity. Based on this observation, one hypothesis to explain the differences between the trees in Figure 4 is that the rearrangements of the tree in Figure 4B occur because the partitions that pass and fail the test differ in their GC content, and that the two trees tend to group together clades with similar GC content (e.g. as in ref (Betancur-r, et al. 2013)). However, it is hard to discern any clear evidence for this from looking at the GC content of the clades presented in figure 4.

**Fig. 3.**
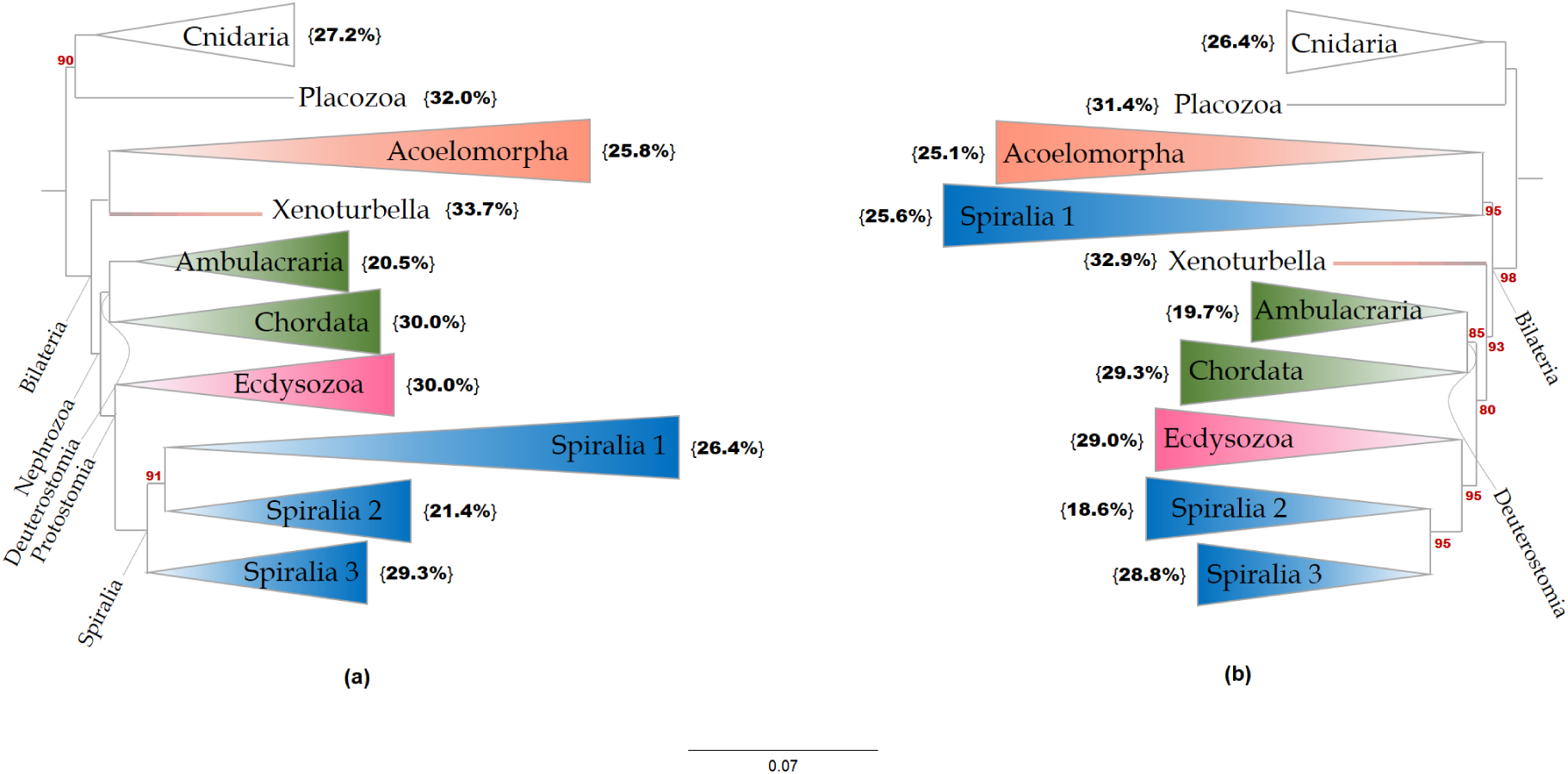
Maximum likelihood trees of Metazoan relationships based on analysis of Cannon 2016 dataset. a) the T_all_ inferred from all 424 partitions and T_pass_ inferred from 281 partitions that passed the MaxSymTest. b) the tree inferred from 143 partitions that failed the MaxSymTest. Red numbers at the internal branches indicate the bootstrap support values that are less than 100% under the best fitting model. Numbers in curly brackets show the GC content of the group. Spiralia 1 consists of Rotifera, Platyhelminthes and Gastrotricha. Spiralia 2 consists of Bryozoa and Entoprocta. Spiralia 3 consists of Annelida, Lophophorata, Nemertea and Mollusca.

The results of our study also provide some insights into the likely cause of model violation in the datasets we examined. Figure 2 shows that violation of marginal symmetry (assessed with MaxSymTest_mar_) was much more common than violation of internal symmetry (assessed with MaxSymTest_int_). This suggests that non-stationarity, which is associated with marginal symmetry, is a more common cause of systematic bias than non-homogeneity in the datasets that we examined (see also Jayaswal, et al. 2005; Ababneh, et al. 2006; Song, et al. 2010). This result hints that the development and application of non-stationary models (e.g. (Yang 1994; Roberts and Yang 1995; Yap and Speed 2005) may be an important avenue towards reducing systematic bias in future analyses. Moreover, our results show a clear preference for simple substitution models with a single transition/transversion ratio over more complex models such as GTR. This suggests that developing non-stationary models with a single parameter for the transition/transversion ratio might be sufficient to reduce systematic bias in phylogenetic analysis.

One limitation of using the tests that we propose in this paper is that their power will be limited if there are few differences between the sequences being examined. Indeed, our analyses show that in our representative sample of more than 3500 partitions from published datasets, roughly ~20% of the variance in whether a partition passes or fails a given test can be attributed to the number of observed differences between the sequences. Nevertheless, this suggests that the remaining ~80% of the variance in whether a partition passes or fails a test could be attributable to other processes, such as variation in the extent of model violation among partitions. This suggests that we should be cautiously optimistic: although a lack of power on small or slowly-evolving partitions may induce some false negatives (i.e. failures to identify partitions that have evolved under non-SRH conditions), the tests we propose still have significant power to identify partitions that show the evidence of model violation. It is possible that removing such partitions from phylogenetic analyses may improve the accuracy of results by reducing the overall burden of model violation on the inference of the tree topology. We hope that our implementation of these tests in user-friendly software IQ-TREE will allow empirical phylogeneticists to continue to explore whether this is the case.

## Acknowledgments

The authors would like to thank Lars Jermiin for discussions and three anonymous referees for providing thoughtful comments on this manuscript.

