## Extended Table for "The Prevalence and Impact of Model Violations in Phylogenetics Analysis"

**Extended Table 1| The proportion of partitions that failed at least one of the three tests - BiSymTest, BiSymTest_mar_, BiSymTest_int_.**

| **Type / genome** | **nuclear** | **mitochondrial** | **plastid** | **virus** |
| --- | --- | --- | --- | --- |
| **1^st^ codon positions** | 49.1% | 63.3% | 33.3% | 87.5% |
| **2^nd^ codon positions** | 42.6% | 30.0% | 33.3% | 50.0% |
| **3^rd^ codon positions** | 85.0% | 80.0% | 33.3% | 87.5% |
| **Other (e.g. intron)** | 33.3% | 100.0% | 83.3% |  |
| **rRNA** | 60.0% | 50.0% |  |  |
| **UCE** | 34.0% |  |  |  |
| **tRNA** |  | 0.0% |  |  |

**Extended Table 2| best-fitting model by ModelFinder and number of partitions that got each model as the best-fit model**. Finding the best-fitting model (which minimize BIC score) for each one of the 3572 partitions.

| **Substitution model** | **#partitions with best-fit model** | **Nucleotide frequencies** |
| --- | --- | --- |
| HKY | 731 | Unequal |
| K80 | 623 | Equal |
| TPM3u | 302 | Unequal |
| TVM | 253 | Unequal |
| TPM2u | 248 | Unequal |
| GTR | 189 | Unequal |
| TPM2 | 152 | Equal |
| TPM3 | 120 | Equal |
| TIM3 | 117 | Unequal |
| K81u | 108 | Unequal |
| TIM2 | 97 | Unequal |
| TN | 89 | Unequal |
| TNe | 82 | Equal |
| JC | 73 | Equal |
| TVMe | 72 | Equal |
| TIM2e | 62 | Equal |
| K81 | 59 | Equal |
| TIM3e | 57 | Equal |
| TIM | 56 | Unequal |
| F81 | 48 | Unequal |
| SYM | 22 | Equal |
| TIMe | 12 | Equal |

**Extended Table 3| best-fitting model by ModelFinder for the partitions that passed each one of the three binomial tests.**

| **BiSymTest** | |  | **BiSymTest_mar_** | |  | **BiSymTest_int_** | |
| --- | --- | --- | --- | --- | --- | --- | --- |
| model | #partitions |  | model | #partitions |  | model | #partitions |
| HKY | 641 |  | HKY | 571 |  | HKY | 702 |
| K80 | 507 |  | K80 | 412 |  | K80 | 597 |
| TPM3u | 257 |  | TPM3u | 201 |  | TPM3u | 295 |
| TPM2u | 215 |  | TPM2u | 178 |  | TPM2u | 240 |
| TVM | 168 |  | TVM | 118 |  | TVM | 238 |
| GTR | 130 |  | TPM2 | 89 |  | GTR | 181 |
| TPM2 | 115 |  | GTR | 73 |  | TPM2 | 145 |
| TPM3 | 105 |  | TPM3 | 71 |  | TPM3 | 116 |
| TIM3 | 91 |  | JC | 69 |  | TIM3 | 113 |
| K3Pu | 77 |  | TIM3 | 64 |  | K3Pu | 103 |
| JC | 73 |  | K3Pu | 59 |  | TIM2 | 95 |
| TN93 | 73 |  | TN93 | 59 |  | TN | 84 |
| TIM2 | 71 |  | TIM2 | 56 |  | TNe | 79 |
| TNe | 65 |  | TNe | 50 |  | JC | 71 |
| F81 | 47 |  | F81 | 43 |  | TVMe | 69 |
| TIM | 41 |  | TIM | 32 |  | K81 | 59 |
| TVMe | 41 |  | TVMe | 31 |  | TIM2e | 57 |
| K81 | 40 |  | K81 | 29 |  | TIM3e | 55 |
| TIM3e | 39 |  | TIM3e | 23 |  | TIM | 52 |
| TIM2e | 34 |  | TIM2e | 21 |  | F81 | 47 |
| SYM | 11 |  | SYM | 9 |  | SYM | 20 |
| TIMe | 7 |  | TIMe | 5 |  | TIMe | 11 |

**Extended Table 4| best-fitting model by ModelFinder for the partitions that passed each one of the three max-value tests.**

| **MaxSymTest** | |  | **MaxSymTest_mar_** | |  | **MaxSymTest_int_** | |
| --- | --- | --- | --- | --- | --- | --- | --- |
| model | #partitions |  | model | #partitions |  | model | #partitions |
| HKY | 492 |  | HKY | 487 |  | HKY | 655 |
| K80 | 363 |  | K80 | 358 |  | K80 | 568 |
| TPM3u | 184 |  | TPM3u | 187 |  | TPM3u | 279 |
| TPM2u | 161 |  | TPM2u | 160 |  | TVM | 240 |
| TVM | 128 |  | TVM | 119 |  | TPM2u | 222 |
| GTR | 114 |  | GTR | 110 |  | GTR | 175 |
| TIM3 | 78 |  | TPM2 | 74 |  | TPM2 | 143 |
| TPM2 | 78 |  | TPM3 | 72 |  | TIM3 | 110 |
| TPM3 | 70 |  | TIM3 | 71 |  | TPM3 | 108 |
| JC | 57 |  | JC | 56 |  | K3Pu | 96 |
| TN | 56 |  | TN | 54 |  | TIM2 | 89 |
| K3Pu | 48 |  | TIM2 | 52 |  | TN | 81 |
| TIM2 | 47 |  | K3Pu | 48 |  | TNe | 75 |
| TNe | 45 |  | TNe | 44 |  | TVMe | 68 |
| F81 | 38 |  | F81 | 39 |  | JC | 63 |
| TIM | 31 |  | TIM | 35 |  | TIM2e | 55 |
| K81 | 29 |  | TVMe | 30 |  | TIM3e | 55 |
| TVMe | 29 |  | TIM3e | 27 |  | K81 | 54 |
| TIM3e | 28 |  | K81 | 26 |  | TIM | 50 |
| TIM2e | 24 |  | TIM2e | 23 |  | F81 | 45 |
| SYM | 10 |  | SYM | 10 |  | SYM | 19 |
| TIMe | 5 |  | TIMe | 7 |  | TIMe | 10 |

**Extended Table 5| best-fitting model by ModelFinder for the partitions that failed each one of the three binomial tests.**

| **BiSymTest** | |  | **BiSymTest_mar_** | |  | **BiSymTest_int_** | |
| --- | --- | --- | --- | --- | --- | --- | --- |
| model | #partitions |  | model | #partitions |  | model | #partitions |
| K80 | 116 |  | K80 | 211 |  | HKY | 29 |
| HKY | 90 |  | HKY | 160 |  | K80 | 26 |
| TVM | 85 |  | TVM | 135 |  | TVM | 15 |
| GTR | 59 |  | GTR | 116 |  | GTR | 8 |
| TPM3u | 45 |  | TPM3u | 101 |  | TPM2u | 8 |
| TPM2 | 37 |  | TPM2u | 70 |  | TPM2 | 7 |
| TPM2u | 33 |  | TPM2 | 63 |  | TPM3u | 7 |
| TVMe | 31 |  | TIM3 | 53 |  | K3Pu | 5 |
| K3Pu | 31 |  | K3Pu | 49 |  | TIM2e | 5 |
| TIM2e | 28 |  | TPM3 | 49 |  | TN | 5 |
| TIM3 | 26 |  | TIM2 | 41 |  | TIM | 4 |
| TIM2 | 26 |  | TIM2e | 41 |  | TIM3 | 4 |
| K81 | 19 |  | TVMe | 41 |  | TPM3 | 4 |
| TIM3e | 18 |  | TIM3e | 34 |  | TNe | 3 |
| TNe | 17 |  | TNe | 32 |  | TVMe | 3 |
| TN | 16 |  | K81 | 30 |  | JC | 2 |
| TIM | 15 |  | TN | 30 |  | SYM | 2 |
| TPM3 | 15 |  | TIM | 24 |  | TIM2 | 2 |
| SYM | 11 |  | SYM | 13 |  | TIM3e | 2 |
| TIMe | 5 |  | TIMe | 7 |  | F81 | 1 |
| F81+I | 1 |  | F81 | 5 |  | TIMe | 1 |
| JC | 0 |  | JC | 4 |  | K81 | 0 |

**Extended Table 6| best-fitting model by ModelFinder for the partitions that failed each one of the three max-value tests.**

| **MaxSymTest** | |  | **MaxSymTest_mar_** | |  | **MaxSymTest_int_** | |
| --- | --- | --- | --- | --- | --- | --- | --- |
| model | #partitions |  | model | #partitions |  | model | #partitions |
| K80 | 260 |  | K80 | 265 |  | HKY | 76 |
| HKY | 239 |  | HKY | 244 |  | K80 | 55 |
| TVM | 125 |  | TVM | 134 |  | TPM2u | 26 |
| TPM3u | 118 |  | TPM3u | 115 |  | TPM3u | 23 |
| TPM2u | 87 |  | TPM2u | 88 |  | GTR | 14 |
| GTR | 75 |  | GTR | 79 |  | TVM | 13 |
| TPM2 | 74 |  | TPM2 | 78 |  | K3Pu | 12 |
| K3Pu | 60 |  | K3Pu | 60 |  | TPM3 | 12 |
| TIM2 | 50 |  | TPM3 | 48 |  | JC | 10 |
| TPM3 | 50 |  | TIM3 | 46 |  | TPM2 | 9 |
| TVMe | 43 |  | TIM2 | 45 |  | TIM2 | 8 |
| TIM3 | 39 |  | TVMe | 42 |  | TN | 8 |
| TIM2e | 38 |  | TIM2e | 39 |  | TIM2e | 7 |
| TNe | 37 |  | TNe | 38 |  | TIM3 | 7 |
| TN | 33 |  | TN | 35 |  | TNe | 7 |
| K81 | 30 |  | K81 | 33 |  | TIM | 6 |
| TIM3e | 29 |  | TIM3e | 30 |  | K81 | 5 |
| TIM | 25 |  | TIM | 21 |  | TVMe | 4 |
| JC | 16 |  | JC | 17 |  | F81 | 3 |
| SYM | 12 |  | SYM | 12 |  | SYM | 3 |
| F81 | 10 |  | F81 | 9 |  | TIM3e | 2 |
| TIMe | 7 |  | TIMe | 5 |  | TIMe | 2 |

**Extended Table 7| The NPD metric mean and confidence interval for BixSymTest, BiSymTest_mar_, and BiSymTest_int_.**

| **BiSymTest** |  | **T_fail_** | **T_pass_** |
| --- | --- | --- | --- |
|  | **T_all_** | 0.40 (0.29,0.52) | 0.67 (0.41,0.94) |
|  | **T_pass_** | 0.82 (0.57,1.06) |  |
| **BiSymTest_mar_** | **T_all_** | 0.31 (0.22,0.40) | 0.65 (0.40,0.90) |
|  | **T_pass_** | 0.72 (0.48,0.96) |  |
| **BiSymTest_int_** | **T_all_** | 0.65 (0.12,1.18) | 0.28 (0.11,0.45) |
|  | **T_pass_** | 0.76 (0.26,1.26) |  |

**Extended Table 8| The proportion of datasets that have a significant p-value in the weighted SH test when using D_pass_ as the input alignment for the test for BixSymTest, BiSymTest_mar_, and BiSymTest_int_.**

|  | **T_all_** | **T_fail_** |
| --- | --- | --- |
| **BiSymTest** | 50% | 87% |
| **BiSymTest_mar_** | 54% | 96% |
| **BiSymTest_int_** | 20% | 90% |

**Extended Table 9| The quartet distances between the three trees (T_all_, T_pass_, T_fail_) in MaxSymTest, MaxSymTest_mar_, and MaxSymTest_int_.**

|  | Dataset | T_all-fail_ | T_all-pass_ | T_fail-pass_ |
| --- | --- | --- | --- | --- |
| MaxSymTest | Anderson_2013 | 942762 | 2630976 | 3479709 |
|  | Bergsten_2013 | 6673 | 18337 | 22924 |
|  | Broughton_2013 | 42756 | 35792 | 43188 |
|  | Brown_2012 | 13784 | 4204 | 17980 |
|  | Cannon_2016a | 6302 | 299483 | 301775 |
|  | Cognato_2001 | 10933 | 2844 | 13777 |
|  | Dornburg_2012 | 9689 | 3608 | 13174 |
|  | Faircloth_2013 | 458 | 156 | 614 |
|  | Lartillot_2012 | 96354 | 0 | 96354 |
|  | McCormack_2013 | 4976 | 6752 | 8828 |
|  | Moyle_2016 | 5751 | 33823 | 38980 |
|  | Oaks_2011 | 10918 | 86433 | 97344 |
|  | Rightmyer_2013 | 596410 | 203550 | 635318 |
|  | Sauquet_2011 | 1347 | 47836 | 48446 |
|  | Wainwright_2012 | 7902579 | 1537295 | 8934454 |
|  | Wood_2012 | 12801 | 2079 | 13750 |
|  | Worobey_2014a | 2681020 | 332059 | 2799987 |
|  | Worobey_2014c | 622591 | 4108 | 624668 |
|  | Worobey_2014d | 111025020 | 16894360 | 123108377 |
|  | Worobey_2014f | 36313021 | 3381394 | 37728429 |
|  | Worobey_2014g | 22846804 | 429497 | 23274926 |
|  | Worobey_2014h | 32055591 | 425390 | 31907851 |
| MaxSymTest_mar_ | Anderson_2013 | 924015 | 2657244 | 3523017 |
|  | Bergsten_2013 | 6661 | 10304 | 15357 |
|  | Broughton_2013 | 40740 | 918 | 41658 |
|  | Brown_2012 | 21778 | 466 | 22060 |
|  | Cannon_2016a | 12836 | 293094 | 305930 |
|  | Cognato_2001 | 11324 | 2844 | 14168 |
|  | Day_2013 | 3644539 | 1661512 | 4521574 |
|  | Dornburg_2012 | 16579 | 1584 | 18004 |
|  | Faircloth_2013 | 458 | 156 | 614 |
|  | Fong_2012 | 433141 | 78033 | 465963 |
|  | Kawahara_2013 | 154971 | 1093 | 155966 |
|  | Lartillot_2012 | 76154 | 0 | 76154 |
|  | McCormack_2013 | 3098 | 10238 | 10622 |
|  | Moyle_2016 | 17193 | 46870 | 64063 |
|  | Oaks_2011 | 30730 | 92387 | 123111 |
|  | Rightmyer_2013 | 532076 | 276570 | 637495 |
|  | Sauquet_2011 | 1347 | 73286 | 73903 |
|  | Wainwright_2012 | 9582787 | 1738223 | 11206752 |
|  | Wood_2012 | 14542 | 2079 | 15214 |
|  | Worobey_2014a | 2677881 | 337644 | 2798877 |
|  | Worobey_2014b | 210251041 | 34373347 | 207469175 |
|  | Worobey_2014c | 258266 | 36220 | 283457 |
|  | Worobey_2014d | 88332246 | 14832614 | 88978592 |
|  | Worobey_2014f | 22033012 | 3417055 | 25040721 |
|  | Worobey_2014g | 25497475 | 429497 | 25925597 |
|  | Worobey_2014h | 33058270 | 427460 | 33460520 |
| MaxSymTest_int_ | Bergsten_2013 | 1130 | 24921 | 25851 |
|  | Cannon_2016a | 3863 | 415440 | 419288 |
|  | Dornburg_2012 | 0 | 54198 | 54198 |
|  | Faircloth_2013 | 0 | 870 | 870 |
|  | McCormack_2013 | 4252 | 5044 | 7704 |
|  | Moyle_2016 | 1368 | 95738 | 95738 |
|  | Oaks_2011 | 0 | 576897 | 576897 |
|  | Rightmyer_2013 | 442526 | 305086 | 606850 |
|  | Wainwright_2012 | 714590 | 23252006 | 23491246 |
|  | Worobey_2014b | 26770407 | 266864710 | 269713210 |
|  | Worobey_2014f | 2826289 | 138623475 | 138564560 |

**Extended Table 10| The quartet distances between the three trees (T_all_, T_pass_, T_fail_) in BiSymTest, BiSymTest_mar_, and BiSymTest_int_.**

|  | Dataset | T_all-fail_ | T_all-pass_ | T_fail-pass_ |
| --- | --- | --- | --- | --- |
| BiSymTest | Anderson_2013 | 2861280 | 914749 | 3668674 |
|  | Bergsten_2013 | 13304 | 7753 | 19467 |
|  | Broughton_2013 | 918 | 42178 | 43096 |
|  | Cannon_2016a | 300248 | 7132 | 307380 |
|  | Cognato_2001 | 7105 | 16536 | 20938 |
|  | Dornburg_2012 | 6679 | 16647 | 22789 |
|  | Faircloth_2013 | 156 | 458 | 614 |
|  | Fong_2012 | 56550 | 448487 | 479314 |
|  | Horn_2014 | 1458294 | 87746 | 1516451 |
|  | Kawahara_2013 | 97177 | 67323 | 135504 |
|  | Lartillot_2012 | 16240 | 103938 | 103938 |
|  | McCormack_2013 | 9649 | 3724 | 7899 |
|  | Moyle_2016 | 93223 | 9929 | 101657 |
|  | Oaks_2011 | 81598 | 32384 | 113822 |
|  | Rightmyer_2013 | 291880 | 461685 | 595840 |
|  | Wainwright_2012 | 1238317 | 6922281 | 7637100 |
|  | Wood_2012 | 2079 | 12997 | 13789 |
|  | Worobey_2014a | 2291125 | 2680004 | 3360983 |
|  | Worobey_2014b | 32497926 | 177369610 | 181427590 |
|  | Worobey_2014d | 16894360 | 112819879 | 126328782 |
|  | Worobey_2014e | 164834729 | 19322469 | 155641064 |
|  | Worobey_2014f | 4264725 | 116898285 | 118315006 |
|  | Worobey_2014g | 105944 | 219155969 | 219177460 |
|  | Worobey_2014h | 441140 | 37772425 | 37899790 |
| BiSymTest_mar_ | Anderson_2013 | 117138 | 4766542 | 4834248 |
|  | Bergsten_2013 | 10446 | 7753 | 16649 |
|  | Broughton_2013 | 35846 | 3026 | 38872 |
|  | Brown_2012 | 754 | 13784 | 14530 |
|  | Cannon_2016a | 7710 | 42191 | 41043 |
|  | Cognato_2001 | 7105 | 16745 | 21148 |
|  | Day_2013 | 1458680 | 3619782 | 4730442 |
|  | Dornburg_2012 | 2285 | 19436 | 21711 |
|  | Faircloth_2013 | 156 | 458 | 614 |
|  | Fong_2012 | 41001 | 415657 | 415894 |
|  | Horn_2014 | 914698 | 323531 | 1143336 |
|  | Kawahara_2013 | 3057 | 144497 | 144766 |
|  | Lartillot_2012 | 0 | 76932 | 76932 |
|  | McCormack_2013 | 5610 | 3891 | 4672 |
|  | Moyle_2016 | 103226 | 37777 | 139224 |
|  | Oaks_2011 | 68502 | 46478 | 114980 |
|  | Rightmyer_2013 | 277643 | 410462 | 529960 |
|  | Wainwright_2012 | 1288727 | 7281784 | 8233317 |
|  | Wood_2012 | 2079 | 13733 | 14644 |
|  | Worobey_2014b | 19098424 | 156325979 | 148609249 |
|  | Worobey_2014e | 161374612 | 29326713 | 150475048 |
|  | Worobey_2014f | 4264725 | 118503641 | 118850210 |
|  | Worobey_2014g | 105944 | 204046143 | 204046560 |
|  | Worobey_2014h | 447720 | 34953228 | 34813762 |
| BiSymTest_int_ | Broughton_2013 | 21367 | 35360 | 56727 |
|  | Cannon_2016a | 613775 | 4332 | 611628 |
|  | Faircloth_2013 | 156 | 0 | 156 |
|  | Horn_2014 | 447482 | 264548 | 694662 |
|  | McCormack_2013 | 14131 | 3272 | 14555 |
|  | Oaks_2011 | 22410 | 13006 | 35407 |
|  | Wood_2012 | 2390 | 6527 | 8813 |
|  | Worobey_2014b | 206405494 | 18975480 | 201655516 |
|  | Worobey_2014f | 26193497 | 4320578 | 30437545 |
|  | Worobey_2014g | 428217 | 22841915 | 23269397 |

**Extended Table 11| Number datasets that contain loci from the different types of genomes and the number of partitions from each type of genome.**

| **Genome type** | **#datasets** | **#genes** | **# partitions** |
| --- | --- | --- | --- |
| Mitochondria | 18 | 30 | 105 |
| Nuclear | 25 | 352 | 3419 |
| Plastid | 2 | 6 | 24 |
| Virus | 8 | 8 | 24 |
