## Extended fig for "The Prevalence and Impact of Model Violations in Phylogenetics Analysis"

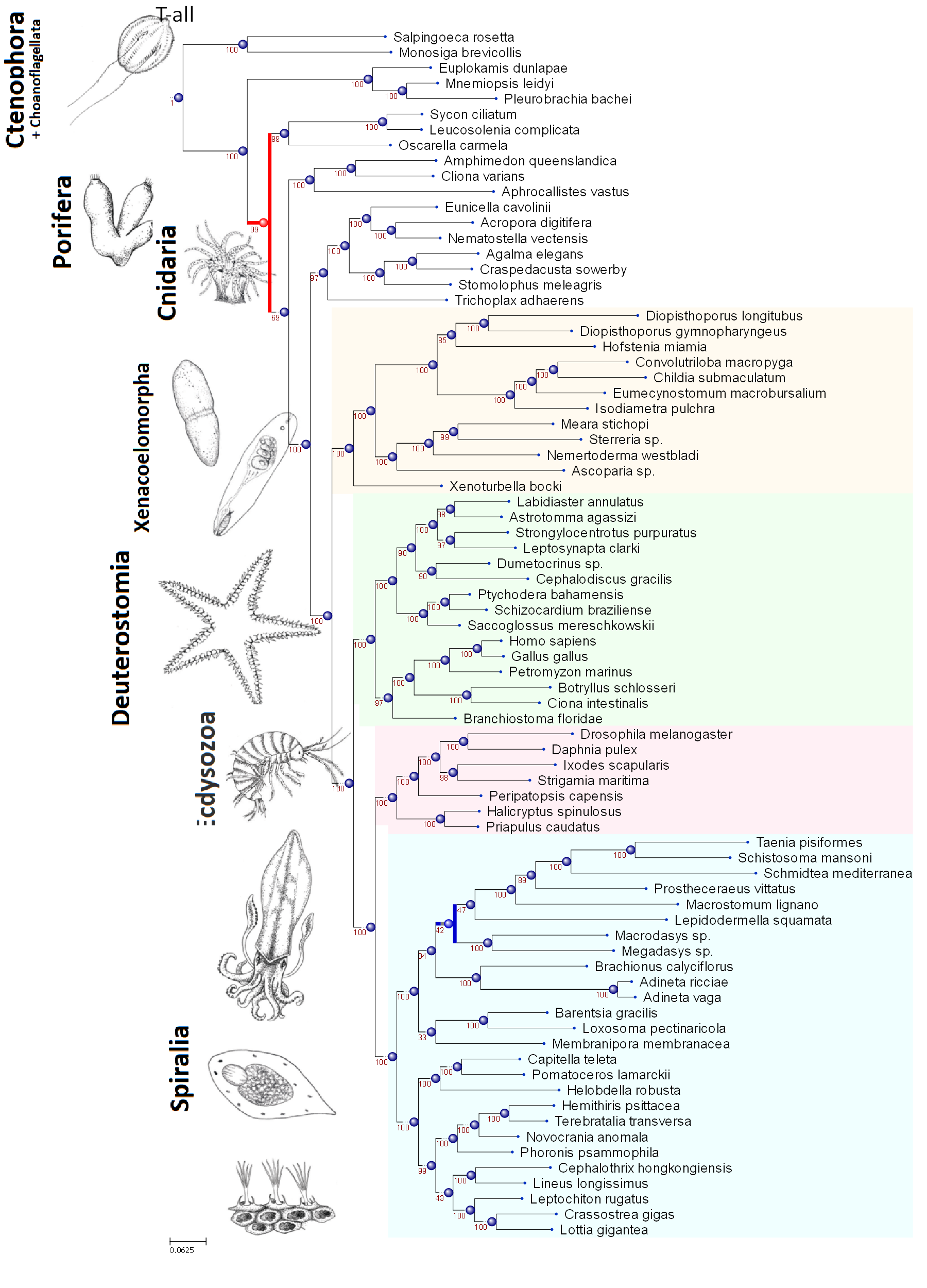

**Extended Figure 1| ML topology of Cannon_2016 dataset inferred from all 424 partitions.**

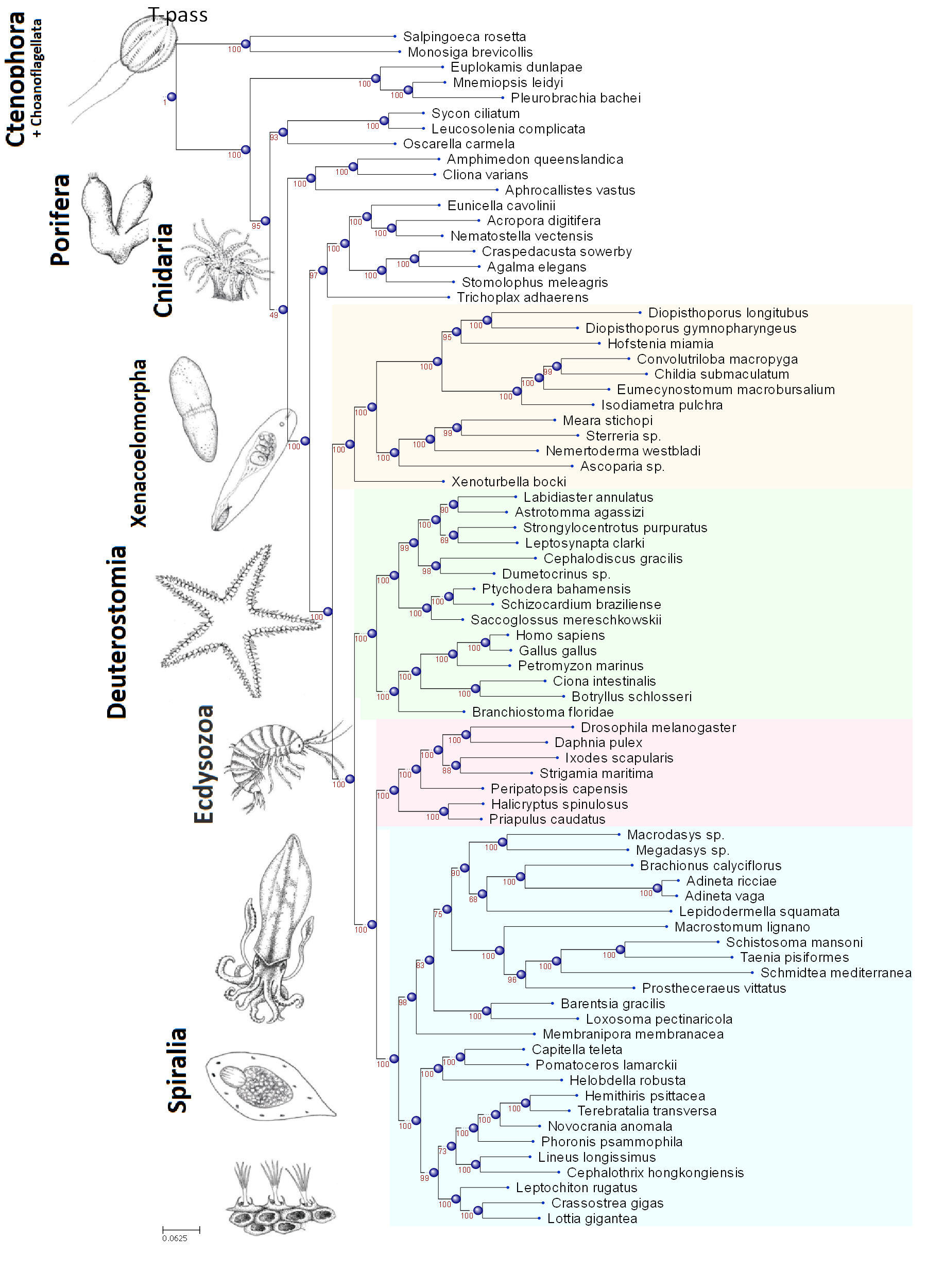

**Extended Figure 2| ML topology of Cannon_2016 dataset inferred from all 342 partitions that passed the BiSymTest.**

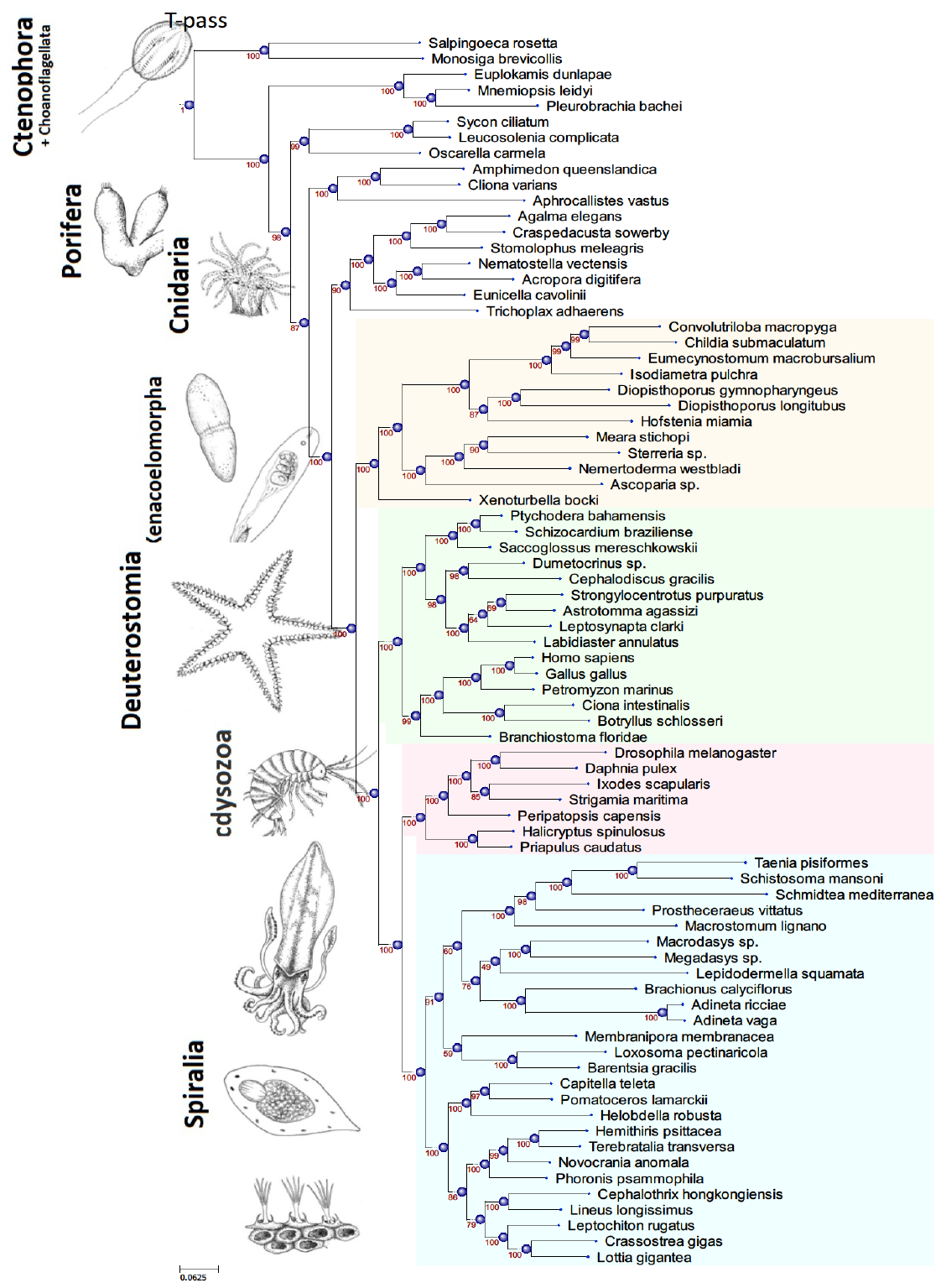

**Extended Figure 3| ML topology of Cannon_2016 dataset inferred from all 281 partitions that passed the MaxSymTest.**

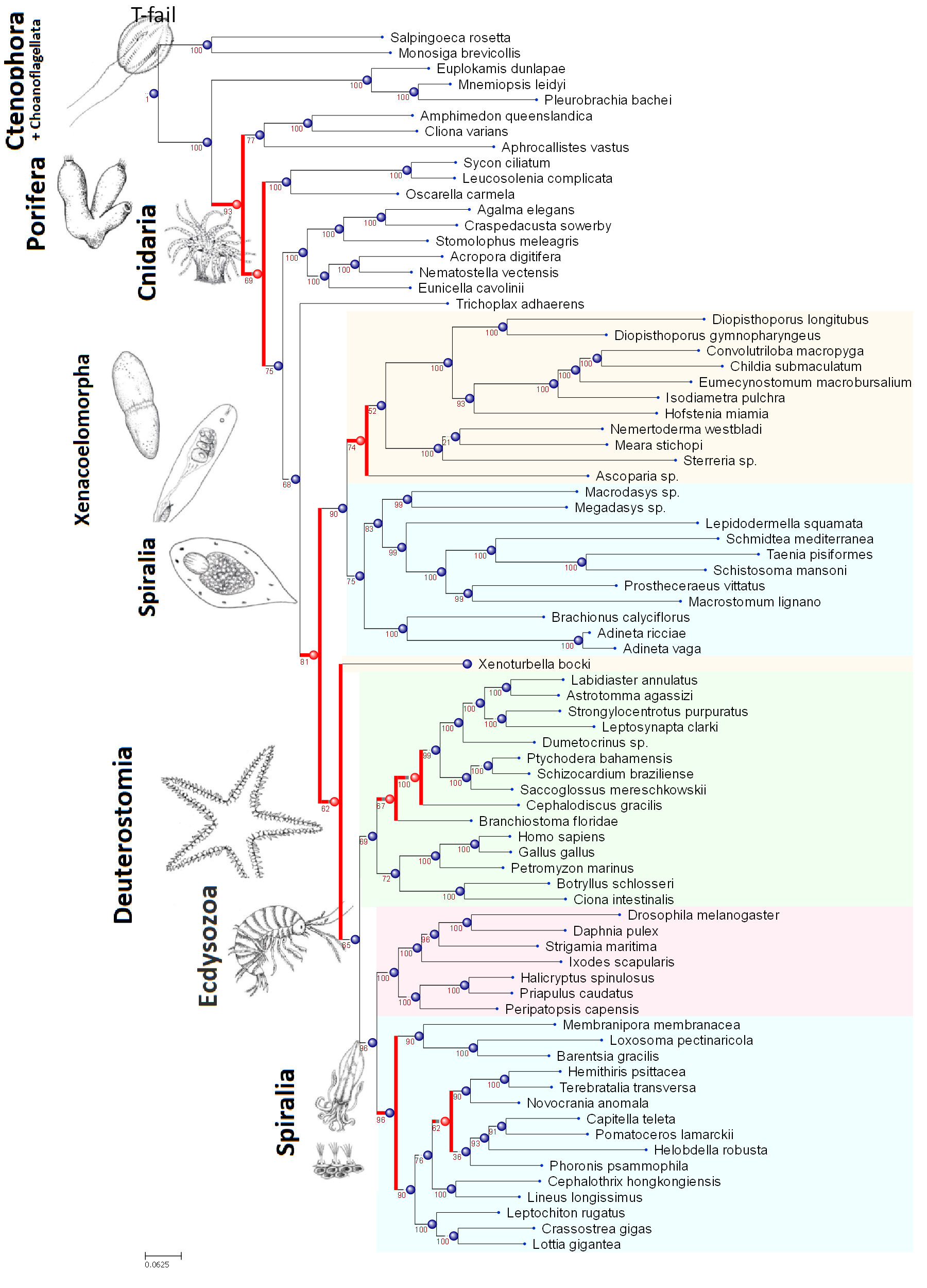

**Extended Figure 4| ML topology of Cannon_2016 dataset inferred from all 82 partitions that failed the BiSymTest.**

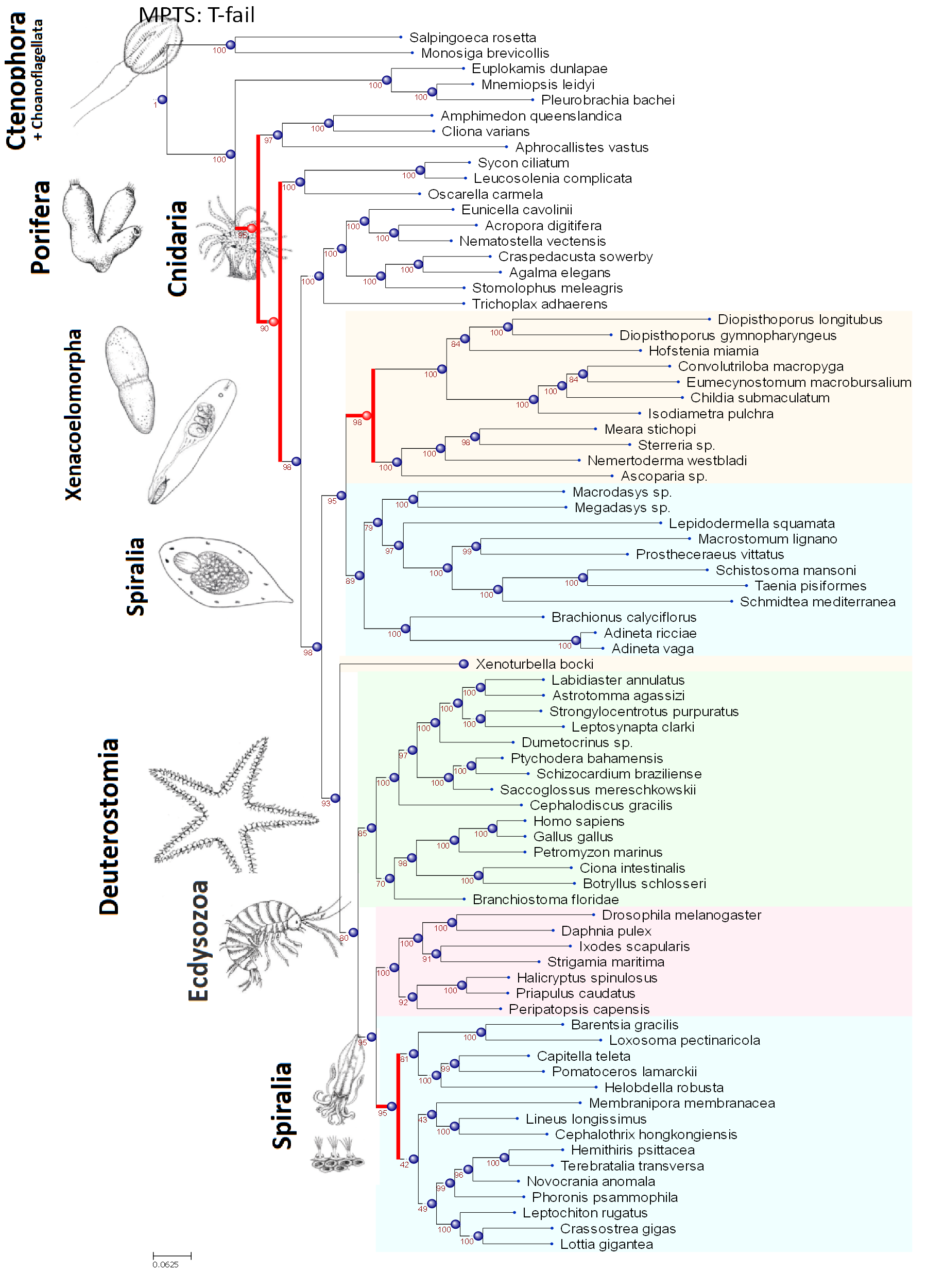

**Extended Figure 5| ML topology of Cannon_2016 dataset inferred from all 143 partitions that failed the MaxSymTest.**

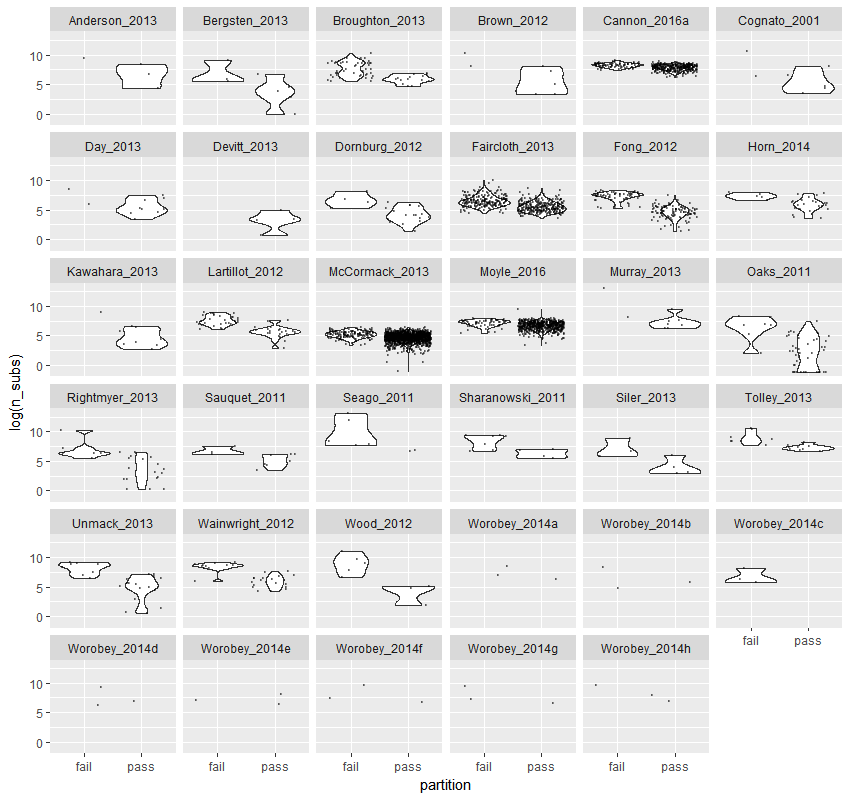

**Extended Figure 6| The number of substitution in partitions that failed or passed the BiSymTest for each dataset.**

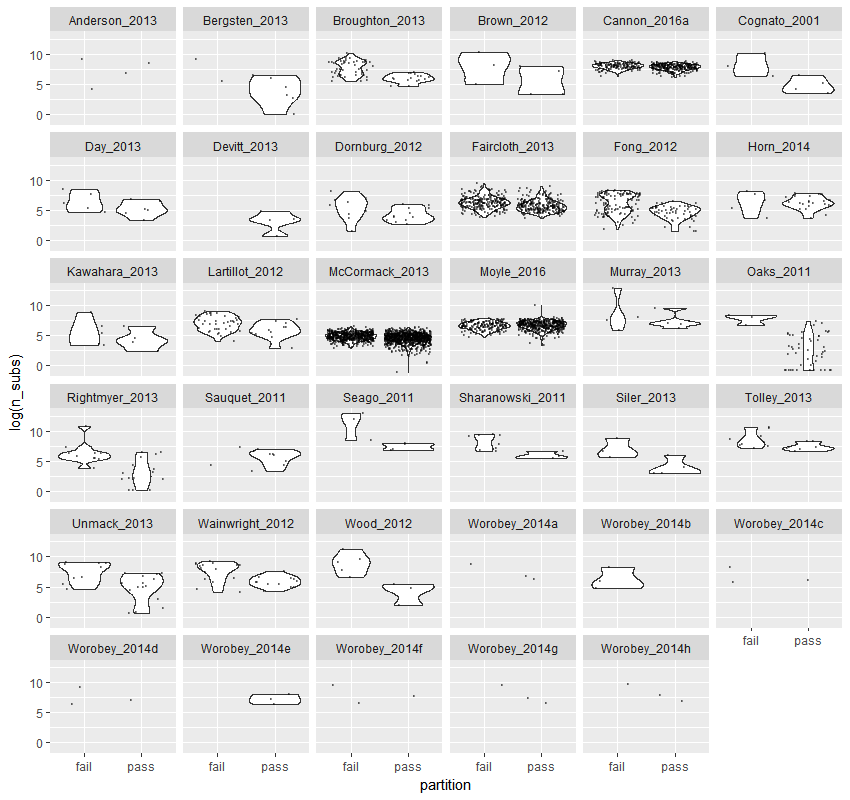

**Extended Figure 7| The number of substitution in partitions that failed or passed the MaxSymTest for each dataset.**

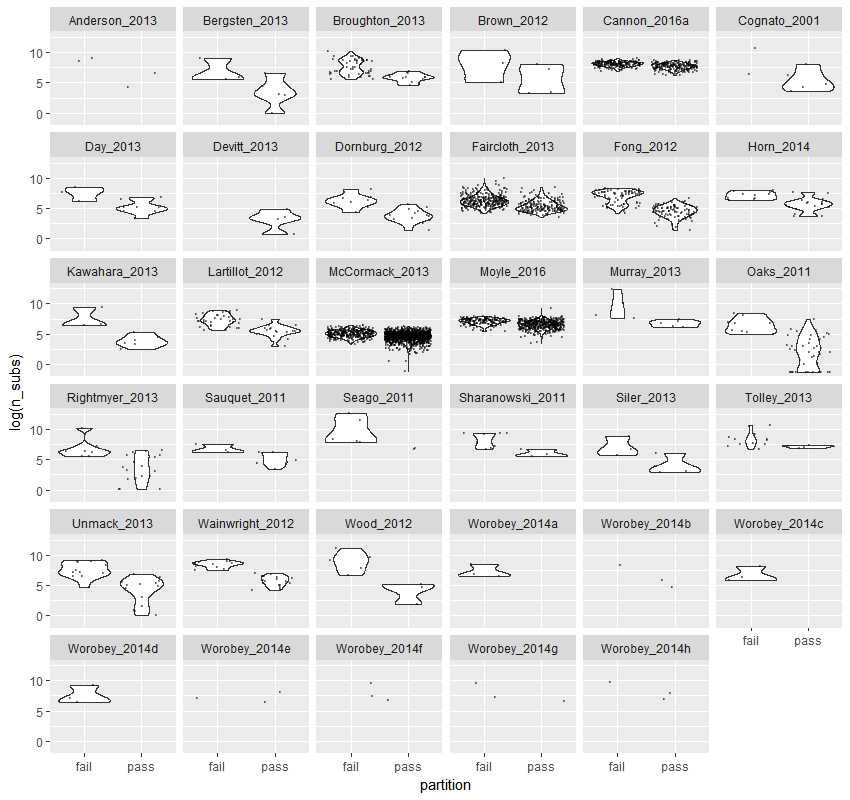

**Extended Figure 8| The number of substitution in partitions that failed or passed the BiSymTest_mar_ for each dataset.**

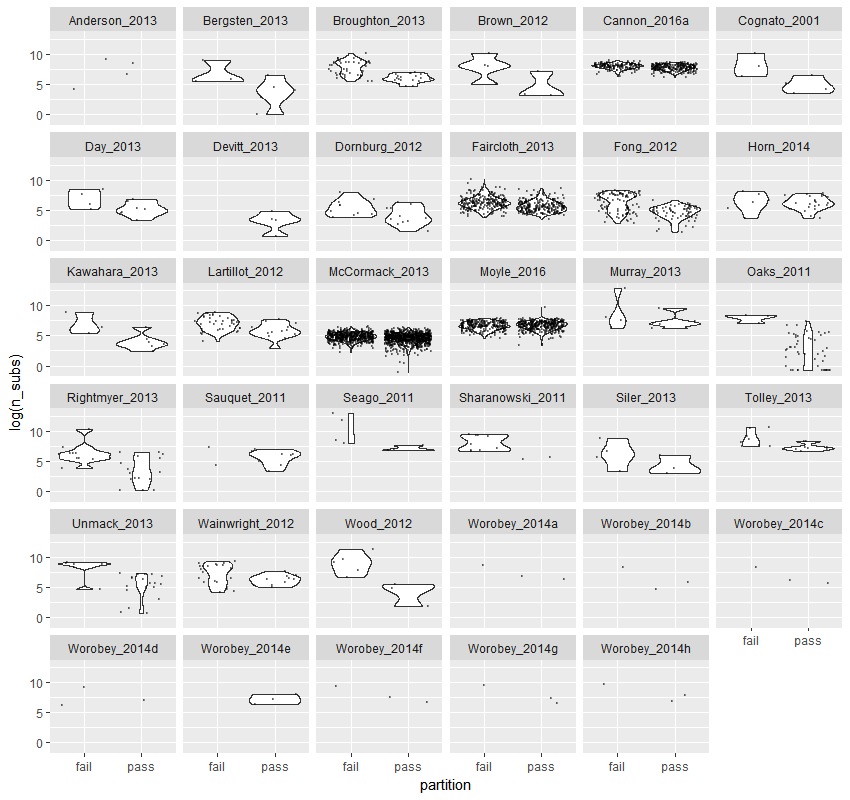

**Extended Figure 9| The number of substitution in partitions that failed or passed the MaxSymTest_mar_ for each dataset.**

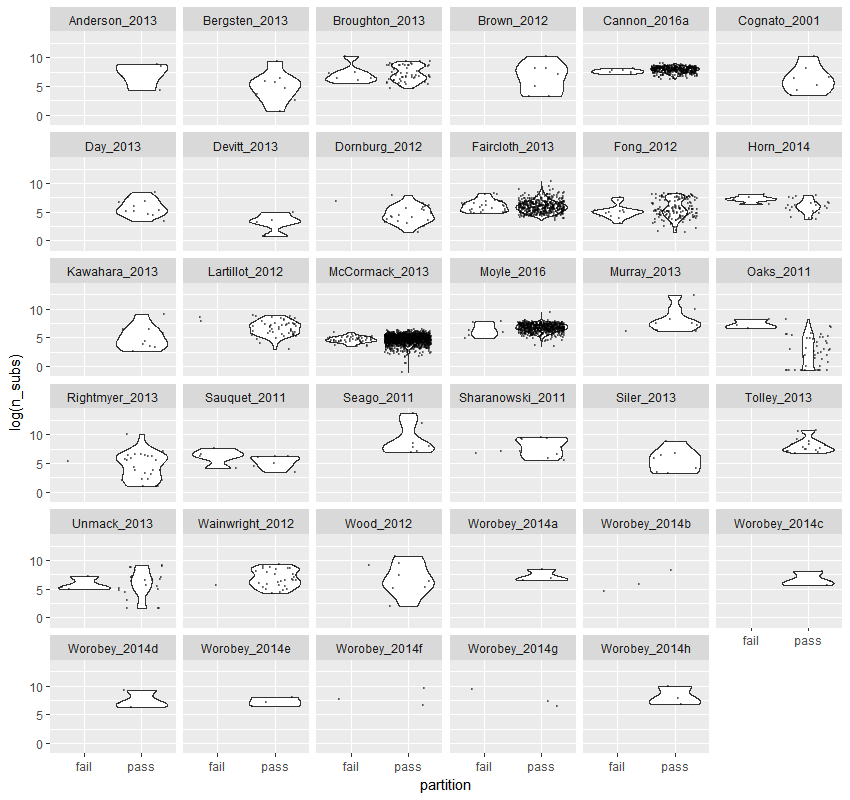

**Extended Figure 10| The number of substitution in partitions that failed or passed the BiSymTest_int_ for each dataset.**

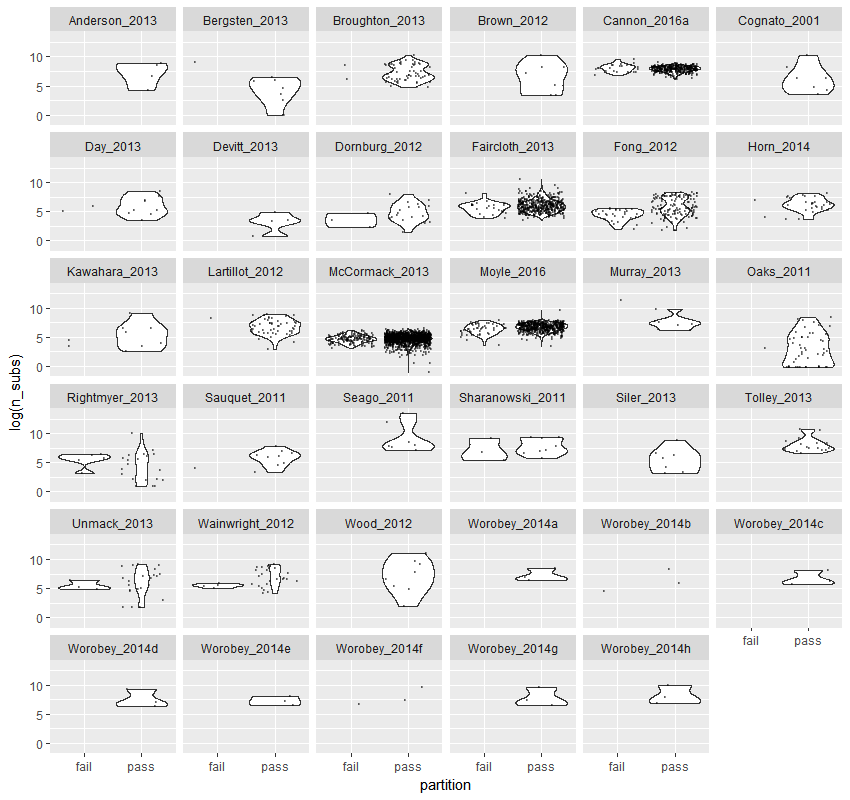

**Extended Figure 11| The number of substitution in partitions that failed or passed the MaxSymTest_int_ for each dataset.**
